# Protocol for capturing the RNA-binding proteome from plants using orthogonal organic phase separation

**DOI:** 10.1101/2025.07.29.667348

**Authors:** Victor A. Sánchez-Camargo, Gertjan Kramer, Harrold van den Burg

## Abstract

RNA-binding proteins (RBPs) regulate many processes related to RNA biogenesis, localization, half-life and function. Here, we present a protocol for the *en masse* isolation of RBPs cross-linked to RNA and provide a strategy for proteomics analysis. We describe the steps for *in vivo* UV-crosslinking of RNA-protein complexes, tissue lysis, fractionation and purification of crosslinked RNA-protein adducts using organic solvents. Although the protocol was developed for *Nicotiana benthamiana* leaves, it can be optimized for use in different plants and tissues.

## Before you begin

RNA-binding proteins (RBPs) regulate RNA-related processes such as RNA splicing, nuclear export, stability, relocalization to e.g., P-bodies, and translation, as well as post-transcriptional gene silencing.^1^ They function by recognizing specific RNA motifs or structures through canonical RNA recognition domains and also through low-complexity regions such as intrinsically disordered domains.^2–4^ Despite their essential roles in development, environmental adaptation, and stress responses, the functional diversity and molecular mechanisms of plant RBPs remain poorly carachterized.^5–8^

Orthogonal Organic Phase Separation (OOPS) is a recently developed method that allows the global identification of RBPs without relying on specific protein tags or polyadenylated RNA.^9,10^ It uses UV light to crosslink RNA-protein complexes *in vivo*, followed by repeated phenol-chloroform extractions, during which RNA-protein adducts accumulate at the interphase (Figure 1). This approach and other variants have been successfully used in animal and bacterial systems, and more recently in plants, where it identified over 2,000 RBPs from Arabidopsis tissues.^10,11^

**Figure 1:**
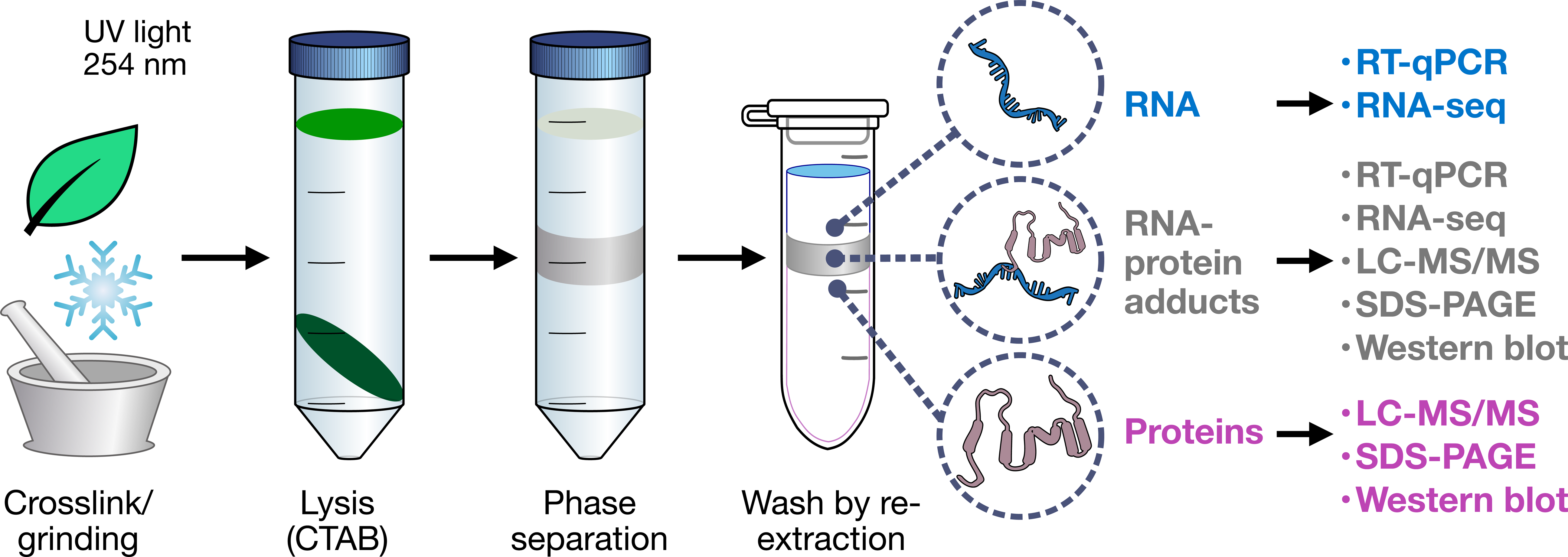
Overview of the OOPS protocol. Plant leaves are harvested and crosslinked using UV light at 254 nm. The tissue is then frozen and ground, and lysis is performed using a custom CTAB buffer, followed by clarification of the lysate. Phase separation with phenol-chloroform produce two main phases (aqueous and organic) and an interphase containing crosslinked RNA-protein complexes. Successive rounds of organic re-extraction remove contaminants from the interphase. High-quality protein and RNA are produced for downstream analysis. RNA and protein drawings were retrieved from NIAID NIH BioArt Source.^25,26^

A major challenge in applying RNA-protein capture methods to plant systems, is the presence of a rigid cell wall, which complicates tissue lysis and limits UV crosslinking efficiency. Additionally, plant tissues can be rich in secondary metabolites, such as phenolics, pigments, and polysaccharides, which interfere with protein extraction and downstream proteomic analysis.^12^

Here, we implemented a modified OOPS protocol for *Nicotiana benthamiana*, incorporating an acidic CTAB (cetyltrimethylammonium bromide)-based lysis step. CTAB is a cationic detergent effective in removing saccharides, compatible with high salt concentrations, and hence high ionic strengths.^13^ Additionally, polyvinylpolypyrrolidone (PVPP), a synthetic polymer, was included to bind polyphenols and other secondary metabolites that may interfere with downstream analyses.^12^ These, in combination with elevated temperatures, create a harsh environment, enhancing the release of organellar content and the disruption of non-specific molecular interactions, thereby improving the purity of the RNA-protein complexes obtained by OOPS.

At the end of this protocol, the user obtains a high-quality preparation of RNA-binding proteins suitable for proteomics analysis (and other applications), as well as the RNA counterpart, which can be used for RT-qPCR and RNA sequencing. Finally, we provide a demonstration of RBP enrichment analysis with proteomics data obtained from *N. benthamiana*.

### Plant growth

**Timing: 5 weeks**

1. Sow *Nicotiana benthamiana* seeds in well-watered potting soil and grow the plants at 25 °C under long day photoperiod conditions (16 h light / 8 dark).

**Note:** A typical experiment includes both UV-crosslinked and non-crosslinked samples. Grow 12 plants per treatment and replicate. We recommend a minimum of four biological replicates per condition.

2. After two weeks, transplant the seedlings into individual pots containing the same soil type, and continue growing them for an additional three weeks.

**Note:** This protocol was developed using the two uppermost fully expanded leaves of 5-week-old plants.

### Key resources table

**CRITICAL:** Ensure that all the reagents employed are of high purity and stored under adequate conditions (refer to vendor’s SDS for safety and handling information).

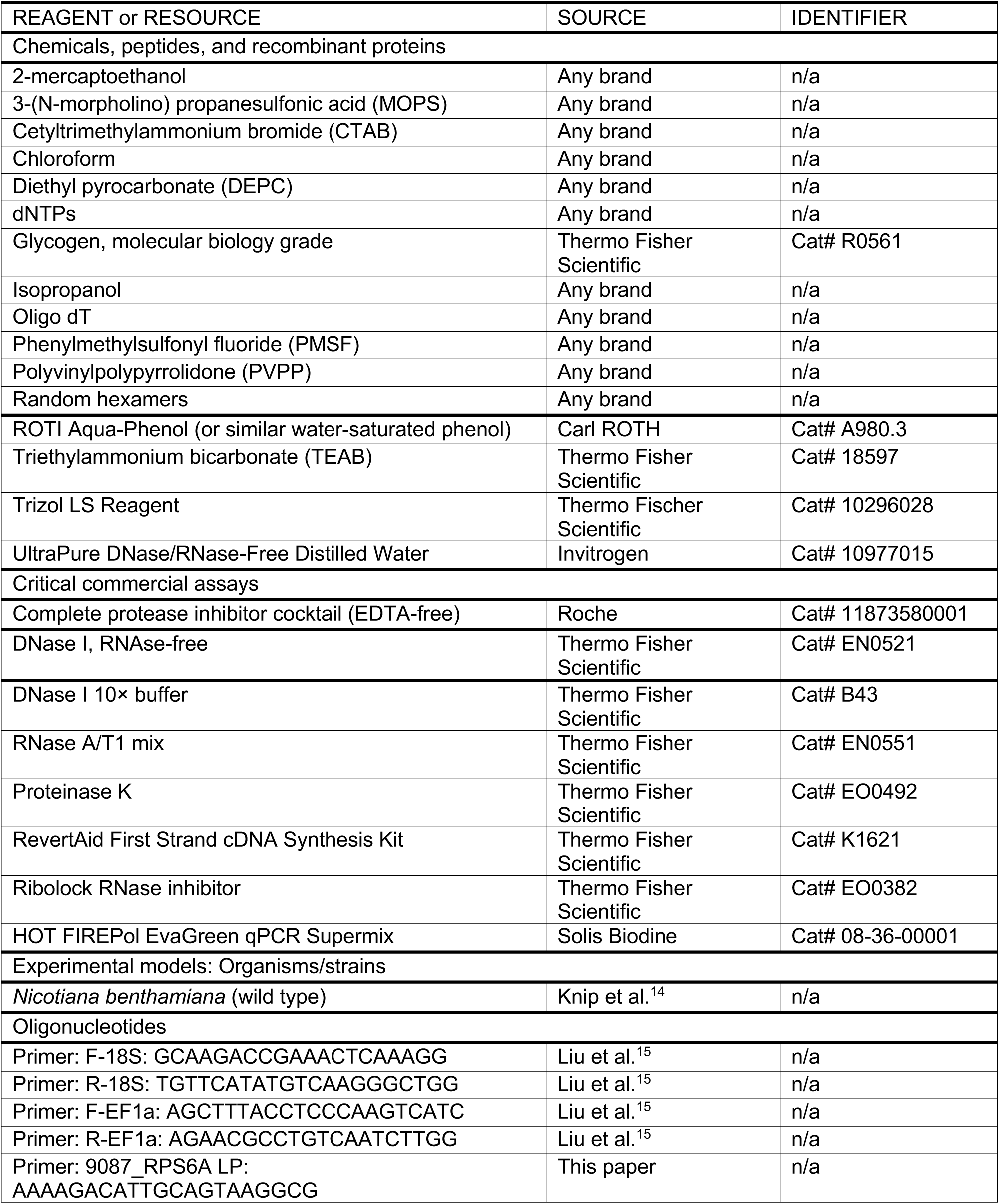
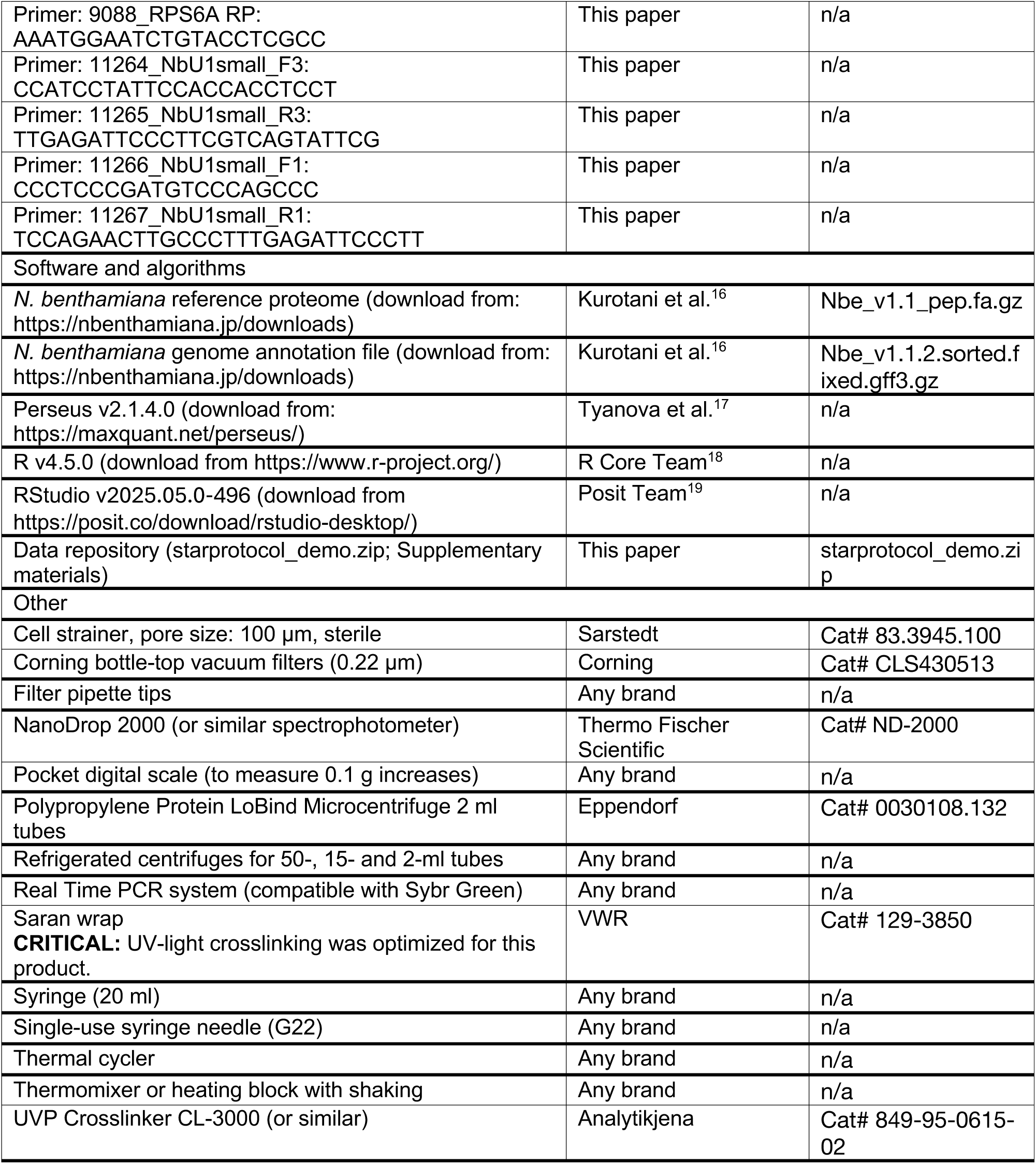

## Materials and equipment setup

### Preparation of buffer stocks

Ensure all the buffers are prepared in advance unless otherwise indicated.

- Lysis buffer (base)

| Reagent | Final concentration |
| --- | --- |
| MOPS | 100 mM |
| CTAB | 2% |
| EDTA | 20 mM |
| NaCl | 1.4 M |

`o` Dissolve all components (in solid form) in deionized water (ddH2O) in a beaker and adjust the pH to 7.0.
`o` Filter using a 0.22 µm bottle-top vacuum filter into a clean, autoclaved bottle containing a magnetic stir bar.

**Optional:** Divide the buffer into aliquots of 250 ml.

`o` Add diethyl pyrocarbonate (DEPC) to a final concentration of 0.2% and mix overnight using a magnetic stirrer.
`o` Remove the magnetic stir bar and autoclave the solution at 121 °C for at least 17 min to inactivate DEPC.

**Note:** DEPC is inactivated at high temperatures, becoming harmless after heat treatment.

**CRITICAL:** MOPS is sensitive to light. To prevent oxidation, protect the container with aluminum foil or use light-opaque glassware.

**CRITICAL:** Lysis buffer (base) is stable for up to 6 months stored at room temperature if protected from light. Do not use the solution if it appears yellowish or contains precipitates.

- DEPC-treated water. Fill 500 ml bottles with ddH2O and add DEPC to a final concentration of 0.1%.
- `o` Mix thoroughly and incubate for at least 2 h at room temperature.
- `o` Sterilize by autoclaving.
- 1 M Tris pH 8.0. Dissolve Tris in DEPC-treated ddH2O to a final concentration of 1 M and adjust the pH to 8.0 using HCl. Sterilize by autoclaving.
- 0.5 M EDTA. Dissolve EDTA in DEPC-treated ddH2O to a final concentration of 0.5 M and adjust to pH 8.0 using NaOH pellets.

**Note:** EDTA is only soluble in water when the pH is adjusted to a basic range, typically around pH 8.0.

5 M NaCl. Dissolve NaCl to a final concentration of 5 M.
`o` Treat with 0.2% DEPC.
`o` Sterilize by autoclaving.
50× Complete protease inhibitor. Dissolve one tablet in 1 ml of 10 mM Tris (pH 8.0). This solution can be stored for up to 12 weeks at −20 °C.
TE-SDS 0.5%. Prepare a solution containing 10 mM Tris (pH 8.0), 1 mM EDTA (pH 8.0) and 0.5 % sodium dodecyl sulfate (SDS) from sterile stocks.
`o` Treat with 0.2% DEPC and mix thoroughly.
`o` Inactivate DEPC by incubating the solution overnight at 37 °C.

**Note:** DEPC treatment is ineffective for concentrated Tris solutions and is only effective at low concentrations.

- 3 M sodium acetate (pH 5.2). Dissolve sodium acetate in ddH2O to a final concentration of 3 M and adjust the pH to 5.2 using glacial acetic acid.
- `o` Treat with 0.2% DEPC and sterilize by autoclaving.
- 0.2 M Sodium citrate buffer (pH 6.0). Dissolve sodium citrate salt to 0.2 M in ddH2O and adjust the pH to 6.0 with HCl.
- `o` Treat with 0.2% DEPC and sterilize by autoclaving.

**CRITICAL:** Store protected from light.

- 70% and 80% ethanol. Dissolve molecular biology-grade ethanol in sterile DEPC-treated water to the desired concentration.
- 2× RNA fragmentation buffer. Prepare a solution containing 20 mM Tris (pH 8.0) and 50 Mm MgCl2 from sterile stocks.
- `o` Treat with 0.2% DEPC and sterilize by autoclaving.
- 100 mM PMSF. Dissolve PMSF in isopropanol to a final concentration of 100 mM. Prepare aliquots of 0.5 ml and store at −20 °C.
- 50 mM TEAB. Prepare the required amount immediately before use from a 1 M stock in UltraPure water.
- 2× Proteinase K reaction buffer.

| Reagent | Final concentration |
| --- | --- |
| Tris (pH8.0) | 50 mM |
| EDTA (pH8.0) | 40 mM |
| SDS | 4% |
| NaCl | 250 mM |

`o` Mix the all the components from sterile stocks.
`o` Treat with 0.2% DEPC and mix well.
`o` Inactivate DEPC by incubating the solution overnight at 37 °C.

### Installation of software and dependencies

- Download and install Perseus (v2.1.3.0) or the latest version.
- `o` Place the extracted folder on your desktop.
- `o` Open the Perseus folder and create a shortcut to the “StartPerseus” executable for quick access and move it to the desktop.

**Note:** Installation on the desktop allows editing Perseus files without administrator privileges.

- Open Perseus and install any required dependencies, following on-screen instructions.

**Note:** This step is only required the first time the program is run.

- Download and install R (v4.5.0) or the latest version.
- `o` **Optional:** Download and install RStudio (v2025.05.0-496) or the latest version.

## Step-by-step method details

### Tissue harvesting and *in vivo* UV-crosslinking

**Timing: variable**

In this step, the plant tissue is harvested and RNA-protein complexes are crosslinked using UV light (254 nm wavelength).

1. Prepare a tray with ice (sized to fit inside the UV-crosslinker) and set the crosslinker’s energy setting to 375 mJ/cm^2^.

**CRITICAL:** The amount of UV light depends on the type of tissue and must be experimentally determined using a dose-response curve. Troubleshooting 1.

2. Harvest the two uppermost leaves from enough plants to obtain 5 g of tissue (typically, 20 leaves are sufficient).
3. Cut a piece of Saran wrap of approximately 50 cm in length. Arrange the leaves with the adaxial surface facing up on the wrap. Fold the wrap to cover and flatten the leaves, forming an envelope (Figure 2).
4. Place the envelope on the ice tray with the adaxial surface facing up. Insert it into the UV-crosslinker and begin irradiation.

**Figure 2:**
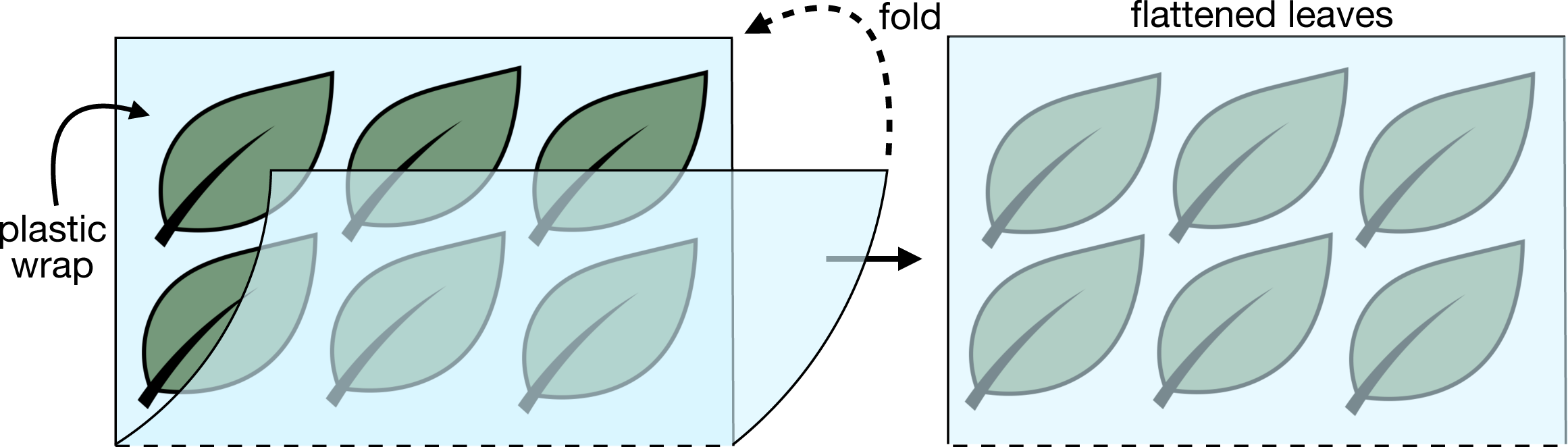
Diagram illustrating the arrangement of leaves for UV crosslinking. Leaves are placed flat on plastic wrap with the adaxial surface facing upward. The wrap is then folded to cover the leaves, ensuring they remain flattened and allowing alternating UV irradiation on both sides.

**CRITICAL:** Ensure that nothing obstructs the UV sensor inside the crosslinker to avoid inconsistencies in UV exposition.

5. Allow the leaves to recover for 30-60 seconds, then flip the envelope and irradiate the abaxial surface. After recovery, irradiate the adaxial surface once more to enhance crosslinking. **Note:** Up to three leaf envelopes can be processed simultaneously by alternating irradiations. The recovery time for one envelope is the irradiation time of the next.
6. Unwrap the envelope and rewrap the leaves with aluminum foil. Label with sample name and date, then snap-freeze in liquid nitrogen. Repeat for all the samples.
7. Grind the frozen tissue to a fine powder and transfer to a sterile 50 ml tube.

**Pause point:** Samples can be stored for at least 3 months at −80 °C until further processed.

### Tissue lysis and phase separation

**Timing: 2.5 h**

In this section, plant tissue is lysed and phenol-chloroform phase separation is performed to isolate the RNA-protein crosslinked complexes from the interphase.

**Note:** All the steps must be conducted in a fume hood to avoid exposure to hazardous solvents.

**CRITICAL:** From this point onwards, all plasticware must be polypropylene to ensure compatibility with phenol and chloroform.

8. Preheat a water bath to 65 °C and chill centrifuges for 50 ml, 15 ml and 2 ml tubes to 4 °C.
9. Prepare the required volume of lysis buffer (approximately 8 ml per sample).
To the Lysis buffer base, add the following to reach final concentrations: 3% PVPP, 1% 2-mercaptoethanol and 1× Complete protease inhibitor cocktail.
Incubate the buffer in a water bath at 65 °C for at least 10 min.
Just before use, add PMSF to a final concentration of 1 mM.

**Note:** This buffer must be freshly prepared. PVPP is insoluble and will make the solution milky. Mix the buffer thoroughly before use, as PVPP precipitates over time.

**CRITICAL:** PMSF is unstable in aqueous solution and should be added immediately before lysis.

10. Weight 5 g of tissue powder into a cold 50 ml tube using a digital pocket scale.

**CRITICAL:** Keep tissue frozen during handling.

**CRITICAL:** This protocol is optimized for 4-5 g or tissue. Using different amounts may require optimization.

11. Allow the tissue to equilibrate at room temperature for 5 minutes inside the fume hood. Add 7.5 ml of pre-warmed lysis buffer and shake vigorously to fully resuspended the tissue.

**CRITICAL:** Manual shaking is more effective than vortexing, as the solution becomes dense and viscous. Adding 4-5 steel beads (⌀4 mm) can enhance tissue homogenizing.

**Note:** A skilled user can process up to eight samples in parallel.

12. Incubate the tubes in a 65 °C water bath for 30 min and shake manually every 5 minutes.

**CRITICAL:** Ensure that tubes remain submerged during incubation to guarantee the temperature.

13. After lysis, cool tubes to room temperature for 5 minutes, then centrifuge at 4,000 × g for 30 minutes at 4 °C to clear the lysate.

**CRITICAL:** If incomplete lysis is observed, adjust the volume of lysis buffer employed in step 11. Troubleshooting 2.

14. Filter the supernatant through a cell strainer (100 µm nylon mesh) into a new 50 ml tube.
15. Collect lysate to prepare input samples (non-crosslinked only).
Transfer 500 µl of lysate from non-crosslinked samples into a 2 ml microcentrifuge tube.
Add 1.5 ml of methanol to precipitate proteins and store at −20 °C until use.
16. To the remaining lysate, add 3 M sodium acetate pH 5.2 to a final concentration of 100 mM. Add one half volume of water-saturated phenol and mix by vortexing for 5 seconds. **CRITICAL:** Use acidic phenol (pH ≤ 6.0) to protect RNA integrity.
17. Add one half volume of chloroform and vortex for 10 seconds. Incubate at room temperature for 5 minutes.
18. Centrifuge at 4,000 × g for 30 min at 4 °C to separate phases.
19. **Optional:** Collect aqueous phase to isolate RNA for quality check and for other downstream applications.
Transfer 1 ml of the aqueous phase (all samples) into a new 2 ml tube for RNA isolation.
Add 120 µl of 5 M NaCl and 880 µl of isopropanol. Mix thoroughly and store at −20 °C. For long-term storage (up to 1 year), keep at −80 °C.
20. Carefully remove and discard the remaining aqueous (top) and organic (bottom) phases using a syringe with a G22 needle. Avoid disturbing the interphase.

**Note:** The interphase contains the crosslinked RNA-protein complexes.

21. Vortex the interphase until homogenized, transfer to a 15 ml centrifuge tube, and centrifuge at 12,000 × g for 5 minutes at 4 °C.
22. Remove all residual aqueous and organic phases. Transfer the interphase to a new 2 ml tube and fill with Trizol LS to the 1.5 ml mark. Vortex thoroughly to resuspend.

**Pause point:** Samples can be stored at −20 °C for up to 1 month before continuing. Use this pause point to synchronize extractions.

### Interphase washing and removal of chromatin traces

**Timing: 2.5 h**

In this section, interphases containing RNA-protein complexes are washed to remove contaminants, including non-RNA-associated proteins and chromatin.

**CRITICAL:** Use filter pipette tips from this point onwards.

23. Before beginning, pre-cool a centrifuge for 2 ml tubes to 4 °C.

**Note:** All centrifugation steps are performed at 4 °C.

24. Thaw the interphases resuspended in Trizol LS (from the step 22) and vortex thoroughly.
25. Add 500 µl of chloroform and shake manually for 30 seconds. Incubate at room temperature for 5 minutes.

**Note:** A skilled user can handle up to 12 samples at a time. Handling more will increase variation in incubation times.

26. Centrifuge 10 minutes at maximum speed (preferably >15,000 × g).
27. Carefully remove and discard the aqueous and organic phases using a syringe (G22 needle), leaving the interphase intact.

**Note:** Work quickly to minimize RNA degradation.

28. Add 150 µl of 1 mM sodium citrate buffer pH 6.0 to the interphase and vortex thoroughly.
29. Add 750 µl of Trizol LS and 250 µl of chloroform. Shake manually for 30 s and incubate for 3 minutes at room temperature.
30. Centrifuge for 3 minutes at maximum speed.
31. Repeat the wash steps (27 to 30) for a total of 3 extractions.

**CRITICAL:** If the interphase becomes diffuse during phase removal, vortex thoroughly and centrifuge again to recompact the material. Troubleshooting 3.

**Note:** Interphases from non-crosslinked samples may appear weak or minimal due to the absence of RNA-protein complexes.

32. After the third wash, discard both the aqueous and organic phases, leaving the interphase intact.
33. Add 300 µl of TE-SDS 0.5% buffer to the interphase and vortex thoroughly.
34. Fill the tubes with methanol and vortex again to disrupt the interphases and precipitate proteins together with associated RNA.
35. Incubate at −20 °C overnight (or for at least 4 h).

**CRITICAL:** Do not store tubes at −80 °C, as this may cause proteins to adhere to the tube walls, reducing recovery.

**Pause point:** Samples can be stored at −20 °C for at least 1 week before proceeding.

36. Centrifuge 30 min at maximum speed at 4 °C to pellet the RNA-associated proteins.
37. Carefully decant the supernatant and wash the pellet with 1 ml of 70% ethanol. Vortex and centrifuge for 5 minutes.
38. Dry the pellet.
Decant the ethanol.
Spin-down the tubes for 30 seconds and use a pipette to remove any remaining ethanol.
Air dry the pellet in a flow hood for exactly 10 min.

**CRITICAL:** Ethanol residues inhibit DNase I activity, while over-drying may make the pellet difficult to resuspend. Troubleshooting 4.

39. Add 305 µl of UltraPure water (see Key Resources Table) and vortex gently until pellet releases. Incubate on ice for 15 minutes to allow hydration.
40. For each sample, prepare a DNase I mix by combining 10 µl of DNase I with 35 µl of 10× DNase buffer in a new tube. Keep on ice.

**CRITICAL:** DNase I is sensitive to mechanical stress. Avoid vigorous pipetting or vortexing.

41. Resuspend the pellets gently using a 1 ml pipette, breaking large fragments into smaller pieces.

**CRITICAL:** Avoid extended pipetting to prevent heat buildup and mechanical RNA fragmentation.

i. 42. Add 45 µl of DNase I mix to each tube and mix gently by flicking the tubes.
ii. 43. Incubate at 37 °C for 1 hour to degrade residual DNA.
iii. 44. Stop the reaction by adding 750 µl of Trizol LS and 250 µl of chloroform. Vortex thoroughly and incubate at room temperature for 3 minutes.
iv. 45. Centrifuge at maximum speed for 3 minutes.

**Note:** At this point, DNase I and degraded nucleic acids partition into the aqueous and organic phases, respectively, while intact RNA-protein complexes remain in the interphase.

46. Carefully remove the aqueous and organic phases, preserving the interphase.
47. Fill the tubes containing the interphase with methanol to precipitate the RNA-protein complexes and remove residual Trizol.
48. Store at −20 °C overnight.

**Pause point:** Samples can be stored at −20 °C for at least 1 week before continuing.

### Elution and purification of RNA-bound proteins

**Timing: 5 h**

In this section, RNA-bound proteins are released by RNA degradation and subsequently precipitated for downstream analysis.

49. Centrifuge the methanol-precipitated samples (from the previous step) at maximum speed for 30 minutes at 4 °C to pellet the complexes.
50. Carefully decant the supernatant. Wash the pellets once with 70% ethanol. Remove residual ethanol and allow the pellets to air dry, as described in previous steps.
51. Resuspend the pellets in 300 µl of UltraPure water. Vortex gently to detach the pellets and allow complete hydration by incubating at 4 °C for 15-30 min, with occasional vortexing. **Note:** During pellet hydration, preheat a heat block to 95 °C.
52. Prepare the samples for RBP release by RNA degradation.
a. Divide the resuspended interphases into two 150 µl aliquots: use one for the following steps and store the other at −20 °C as a backup.

**CRITICAL:** If RNA will be analyzed (e.g., RT-qPCR or RNA-seq), use the backup sample for RNA purification (steps 84a-i to 89).

b. Add 150 µl of 2× RNA fragmentation buffer and mix thoroughly.
53. Incubate at 95 °C for 7 minutes in a heat block to fragment the RNA chemically.
54. Cool down the samples to room temperature for 2 min, then add 10 µl of RNase A/T1 mix and vortex thoroughly.

**CRITICAL:** RNase A/T1 will remain in the sample and it is typically among the most abundant proteins detected by LC-MS/MS. Do not increase the enzyme concentration. Troubleshooting 5.

55. Incubate at 37 °C for 3 hours to complete RNA degradation.
56. Stop the reaction by adding 300 µl of Trizol LS and 100 µl of chloroform. Vortex thoroughly and incubate for 3 minutes at room temperature.
57. Centrifuge at maximum speed for 5 minutes to separate phases.
58. Carefully tilt the tube and use a 200 µl pipette to extract 190 µl from the bottom of the organic phase. This contains the proteins released from RNA after digestion.

**CRITICAL:** Tilting the tube displaces the interphase, allowing more precise access to the organic phase. Avoid contaminating the sample with interphase, which may carry glycoproteins and other impurities.

59. **CRITICAL:** Transfer the organic phase into a polypropylene Protein LoBind 2 ml tube.
60. To precipitate the released RBPs, add 1.8 ml methanol and incubate overnight at −20 °C.

**Pause point:** Samples can be stored at −20 °C for up to one week prior to further processing.

### Resuspension of (RNA-binding) proteins

**Timing: 1.5 h**

In this section, proteins from both input samples and interphases are cleaned, concentrated, and prepared for downstream analysis such as SDS-PAGE, western blot, or LC-MS/MS.

61. Centrifuge the protein input samples (from step 15) and the precipitated RBPs (from step 60) at maximum speed for 30 min at 4 °C to pellet the proteins.
62. Wash the pellets once with 80% ethanol, followed by a second wash with 70% ethanol. Decant the ethanol and allow the pellets to air dry as described earlier.

**Note:** Residual Trizol in the samples may lead to overestimation of protein concentration in colorimetric assays such as BCA and Bradford.

63. Resuspend the input pellets in 800 µl of 50 mM TEAB (or buffer of choice) and the RBP pellets in 50 µl. Allow the pellets to hydrate at 42 °C for 10 minutes. If possible, use a thermomixer or heat block with shaking at >1000 rpm to facilitate resuspension.

**Note:** For LC-MS/MS analysis, we recommend resuspending the proteins in 50 mM triethylammonium bicarbonate (TEAB). Confirm the required buffer composition with your proteomics service provider.

64. Prepare samples for LC MS/MS analysis following the instructions of your provider.

### Analysis of proteomics data

**Timing: 1.5 h**

In this section, the mass spectrometry data generated using the samples produced at the end of the previous section, will be analyzed using Perseus software^17^ to identify RBPs enriched in crosslinked fractions. After running the provided R scrip, the user will generate two volcano plots comparing the enriched proteins before and after filtering.

**Note:** Mass spectra are first processed with MaxQuant^20^ to perform peptide-spectrum matching and quantification. The *N. benthamiana* reference proteome^16^ (see Key Resources Table) is used for protein identification. Confirm with your proteomics provider the type and format of output files you will receive.

**CRITICAL:** For the demonstration script to run without modifications, retain the file names provided in the supplementary materials and in this section.

65. After peptide-spectrum matching, a tabular file containing protein groups will be generated.
66. Download the provided data package (starprotocol_demo.zip) from the Key Resources Table and extract it in your desired working directory.
67. Load the *N. benthamiana* annotation file on Perseus.
Navigate to: Desktop > Perseus > bin > conf > annotations.
Copy the file annot_nbenth_MPP_v7.txt into this directory

**Note:** The annotation file is a tab-separated values file and can be explored using spreadsheet software (e.g., Microsoft Excel).

**CRITICAL:** Ensure that the protein identifiers in your annotation and sequence database are compatible. If using custom proteomes, annotate them using tools like eggNOG-mapper.^21^

68. Load the proteinGroups.txt file in Perseus:
Go to: Matrix > Generic matrix upload.
Select the columns containing the LFQ values from the list (18 columns labeled “LFQ intensity…”).
Move them to the “Main” section pressing the “>” button.
Press OK to generate the matrix for analysis.
69. Clean-up the matrix:
Remove the following rows using: Filter rows > Filter rows based on categorical column.
Potential contaminants.
Reverse hits.
Proteins identified only by site.

**Note:** In the provided dataset, this will reduce the example data from 4309 to 4068 entries.

70. Group the biological replicates by sample type:
Go to: Annot. rows > Categorical annotation rows.
Select: “Read from file” and choose the provided file grouping_fraction.txt.

**Note:** Alternatively, annotations can be done manually by selecting replicates and assigning a group name.

71. Filter the identified proteins for reproducibility:
Apply: Filter rows > Filter rows based on valid values.
Select the option “Percentage” and set it to 70.
Select “In at least one group”, to keep only proteins present in ≥5/6 replicates, in at least one fraction.
72. Add annotations to the identified protein groups:
Go to: Annot. columns > Add annotation.
Select the annot_nbenth_MPP_v7.txt file and import desired metadata. For demo purposes, load all the listed annotations.
73. Log2 transform the LFQ data.
Apply: Basic > Transform.
74. Impute missing values to allow statistical comparisons:
Apply: Imputation > Replace missing values from normal distribution. Use default settings.

**Note:** The matrix generated at this point can be analyzed using the software of preference (e.g., R).

75. Perform statistical testing to identify enriched proteins:
Go to: Tests > Two-samples test.
Assign “OOPS_XL” to the group 1 (top) and “OOPS_NOXL” to the group 2 (bottom).
Select “Welch’s T-test”.

**CRITICAL:** A non-parametric test is employed due to missing values in non-crosslinked samples.

76. Export the results of the Welch’s T-test.
Go to: Export > Generic matrix export.
Save the new matrix as starprotocol_rbpome.txt.
77. Filtering enriched RBPs.
Open starprotocol_rbpome.txt in a spreadsheet program.
**CRITICAL:** Delete the second row (Perseus metadata), which can interfere with downstream analysis.
Filter the column “Welch’s T-test Significant OOPS_XL_OOPS_NOXL” to retain only “+” entries. This will reduce the dataset from 4068 to 431 protein groups.
Apply a second filter on “Welch’s T-test Difference OOPS_XL_OOPS_NOXL” to retain proteins with a fold change ≥1. This yields 369 significantly enriched RBPs.
Save the filtered list as starprotocol_enriched_RBPome.txt.
78. **Optional:** Visualization of enriched RBPs (volcano plot).
Ensure that both the starting matrix (starprotocol_rbpome.txt) and the filtered matrix (starprotocol_enriched_RBPome.txt) are in the same folder.
Run R on the terminal or open RStudio.
Set the provided script volcano_rbpome.R.
Update the section “## 0. Set working directory” with the location of your files.
Save changes.
Run the script to generate a volcano plot.
A new figure will be stored in your working folder (volcano_rbps_starprotocol.pdf) showing data before and after filtering.

**Note:** If working with custom data, update the file names in section “## 2. File paths of the script” accordingly:

file_unf <-“starprotocol_rbpome.txt”

file_flt <-“starprotocol_enriched_RBPome.txt”

79. Use the list of enriched RBPs (step 77) for further analyses, validations or hypothesis generation.

CRITICAL: Perform quality checks on the mass spectrometry data to ensure the enriched proteins are true RBPs (see Expected outcomes section). Troubleshooting 6.

### Purification of high-quality RNA with DNase and proteinase K

**Timing: variable**

In this section, high-quality RNA is isolated for downstream applications such as electrophoresis, RT-qPCR or RNA-seq.

80. Centrifuge the RNA-containing samples collected in step 19 for 15 min at max speed at 4 °C to pellet the RNA.
81. Wash the resulting RNA pellet with 1 ml of 70% ethanol and vortex thoroughly.
82. Centrifuge for 5 minutes at maximum speed. Remove any residual ethanol and air dry the pellet as previously described.
83. Decant the supernatant and resuspend the pellet in 500 µl of 1 mM sodium citrate (pH 6.0) and quantify the RNA using a NanoDrop or equivalent spectrophotometer.

**Note:** A260/A280 ratios are expected to be <2.0 due to proteins covalently bound to the RNA in crosslinked fractions. A260/A230 should be ∽2.2, indicating minimal salt and solvent contamination.

84. Purification of RNA with DNase and proteinase K.
Transfer 30-50 µg of RNA to a new 2 ml tube. Store the remaining sample at −80 °C.
**OPTIONAL:** If using the interphase containing RNA-protein adducts (backups from step 52a), quantify the RNA using a NanoDrop and transfer 30-50 µg to a new 2 ml tube.
**OPTIONAL:** For non-crosslinked interphase controls, use the same volume as the corresponding crosslinked fraction (these often yield minimal RNA).
Adjust all sample volumes to 85 µl with UltraPure water.

**Note:** Keep sample volumes minimal to maintain concentration.

c. Prepare DNase I master mix by adding 5 µl of DNase I and 10 µl of 10× DNase buffer per sample. Mix gently.
d. Add 15 µl of DNase I master mix to each RNA sample and mix gently by pipetting.
e. Incubate at 37 °C for 45 min.
f. To stop the reaction, add 110 µl of 2× proteinase K reaction buffer.
g. Add 10 µl of proteinase K to each sample and mix thoroughly.
h. Incubate at 55 °C for 1.5 h (use shaking at 550 rpm if available) to degrade the DNase and crosslinked proteins on the RNA.
85. After digestion, add 660 µl of TE-SDS 0.5%, 120 µl of 5 M NaCl, 2 µl of glycogen. Vortex thoroughly.
86. Add 1 ml of isopropanol and mix thoroughly. Incubate overnight at −20 °C to precipitate the purified RNA.
87. Centrifuge at maximum speed for 30 minutes at 4 °C and remove the supernatant.
88. Wash the pellet twice with 70% ethanol and dry the pellet as before.
89. Resuspend the RNA in 50 µl of 1 mM sodium citrate (pH 6.0) and quantify by your preferred method.

### RNA analysis by RT-qPCR

**Timing: 5 h**

In this section, purified total RNA and deproteinated RNA are reverse transcribed into cDNA and analyzed via quantitative PCR to detect transcript abundance.

90. Reverse transcription (cDNA synthesis).
Prepare the following mix to anneal primers for reverse transcription. Mix for RNA priming

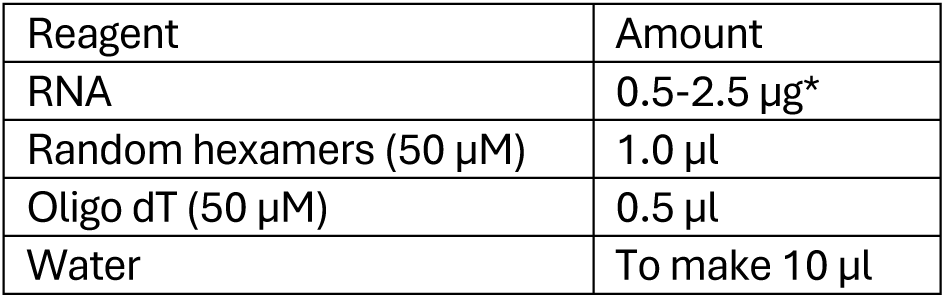

| Reagent | Amount |
| --- | --- |
| RNA | 0.5-2.5 μg* |
| Random hexamers (50 μM) | 1.0 μl |
| Oligo dT (50 μM) | 0.5 μl |
| Water | To make 10 μl |

**\*CRITICAL:** For non-crosslinked interphase controls, use the same volume as the corresponding crosslinked fraction (the RNA yield in these samples is minimal).

**CRITICAL:** Random hexamers are required because this protocol isolates both polyadenylated and non-polyadenylated RNAs.

b. Denature RNA-primer mix at 70 °C for 5 min in a thermal cycler.
Immediately transfer the tubes to ice to stabilize primer annealing.
91. Prepare a reverse transcription (RT) master mix: Reaction mix for RT

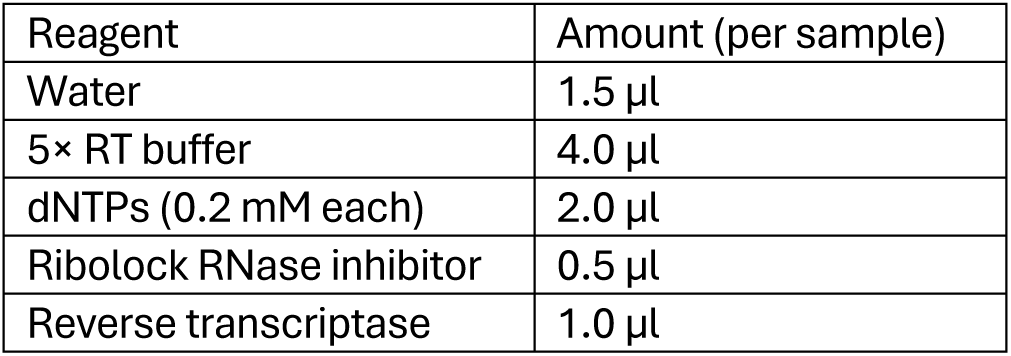

| Reagent | Amount (per sample) |
| --- | --- |
| Water | 1.5 μl |
| 5× RT buffer | 4.0 μl |
| dNTPs (0.2 mM each) | 2.0 μl |
| Ribolock RNase inhibitor | 0.5 μl |
| Reverse transcriptase | 1.0 μl |

RT program

Add 9 µl of RT master mix to each RNA-primer mix. Mix thoroughly and briefly spin down.
Run the RT reaction in a thermal cycler using the following program:

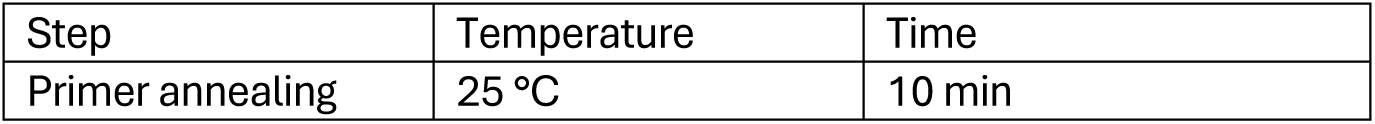
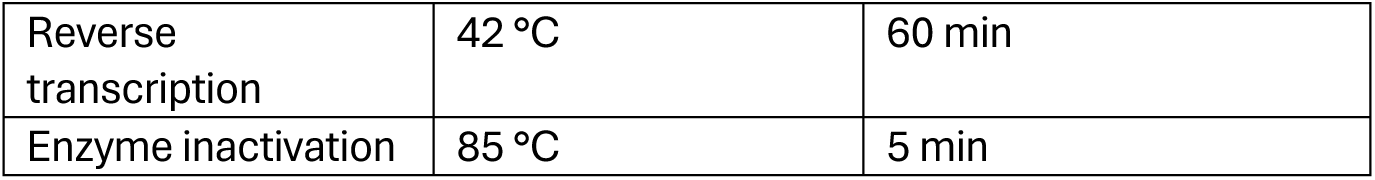

c. After the RT reaction, adjust the cDNA concentration to 25 ng/µl (of input RNA) using 10 mM Tris pH (8.0).

**Pause point:** Store the cDNA at −20 °C or −80 °C for long-term until use.

92. Perform quantitative PCR using cDNA.
Prepare a PCR master mix as follows (in this order): qPCR master mix

| Reagent | Amount (per sample) |
| --- | --- |
| Water | 9.0 μl |
| HOT FIREPol EvaGreen qPCR Supermix (5×) | 3.0 μl |
| Primer 1 (3 μM) | 1.0 μl |
| Primer 2 (3 μM) | 1.0 μl |

b. Add 14 µl of qPCR master mix to the corresponding tubes or plates.
c. Add 1.0 µl of cDNA to each tube/well.
d. Spin down for 1 min at 1000 × g at room temperature.
e. Run real time PCR using the following cycling conditions: qPCR cycling protocol

| Step | Temperature | Time | Cycles |
| --- | --- | --- | --- |
| Initial activation | 95 °C | 12 min | 1 |
| Denaturation | 95 °C | 15 sec | 40 cycles |
| Annealing/extension | 60 °C (detection in Sybr Green channel) | 60 sec |  |
| Melt curve | n/a | n/a | n/a |

**CRITICAL:** Perform a standard curve to determine primer efficiency, optimal cDNA concentration, and presence of PCR inhibitors (e.g., residual phenol or alcohol).

93. Calculate relative transcript abundance in the different fractions using the ΔCt method. Normalize Ct values to the non-crosslinked aqueous phase, which serves as the reference (=1) (Figure 3). Use the following formula:

**Figure 3:**
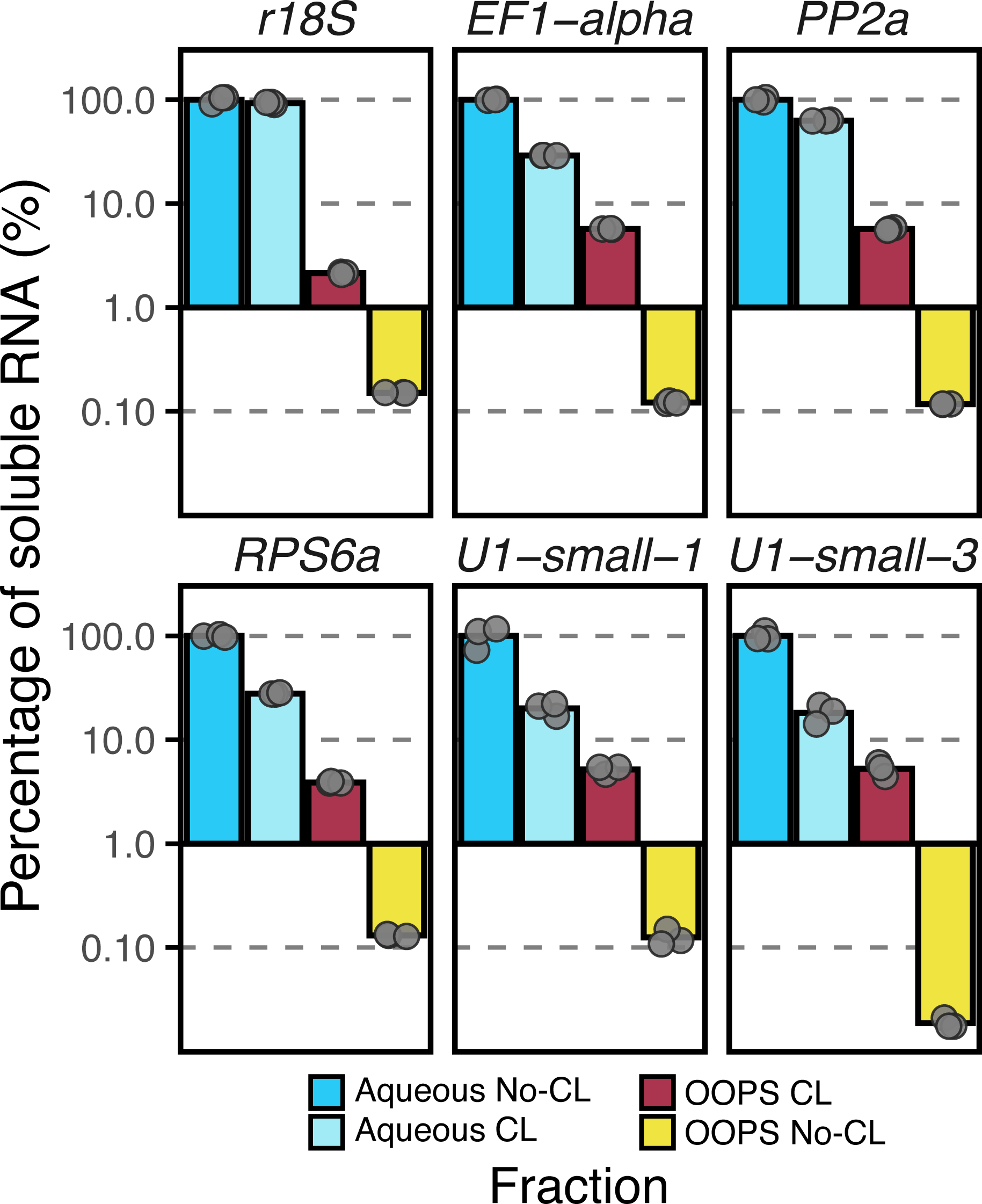
RT-qPCR analysis of RNA species across different fractions. R18S (large ribosomal subunit); EF1-alpha, PP2a, and RPS6a (mRNAs); and U1-small-1 and U1-small-U3 (snoRNAs) were analyzed. The highest RNA levels are detected in the aqueous fraction of non-crosslinked (No-CL) samples, which reflects the total transcriptome. Following crosslinking, the RNA bound to proteins is redistributed to the interphase, resulting in a moderate decrease in RNA abundance. In crosslinked (CL) samples, the RNA-protein complexes accumulate in the interphase. In the absence of UV exposure, RNA levels in the interphase remain negligible.

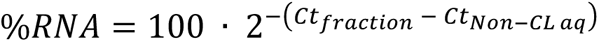

Variables

*Ct_fraction_* = averaged Ct values obtained using RNA from a given fraction

*Ct_Non-CL aq_* = averaged Ct values obtained using RNA from the aqueous phase of non-crosslinked samples

**Note:** If different amounts of input tissue or RNA for cDNA synthesis are used, values must be corrected accordingly.

**CRITICAL:** Internal normalization genes (e.g., housekeeping genes) cannot be used in this protocol due to the nature of in vivo UV crosslinking and fractionation-specific RNA enrichments. Expression comparisons are made only between fractions, not across genes or treatments.

### Expected outcomes

To determine the optimal UV crosslinking conditions for *N. benthamiana* leaves, we performed a dose-response analysis. As shown in Figure 4, we applied incremental doses of UV light (75 mJ/cm^2^ per step) to the leaves, followed by sample fractionation using the OOPS protocol. RNA and proteins were then isolated from the aqueous and interphase (OOPS) fractions, respectively, and quantified. We observed that RNA yield in the aqueous phase progressively decreased with increasing UV doses. Conversely, both RNA and protein levels in the interphase increased, suggesting reduced RNA solubility, likely due to the stabilization of RNA-protein complexes upon UV light exposure, which resulted in their partitioning into the interphase. At the highest UV dose tested, we observed a decrease in RNA yield in the OOPS fraction, indicating possible degradation or over-crosslinking. Therefore, 375 mJ/cm^2^ was determined to be the optimal condition for efficient *in vivo* crosslinking of RNA-protein complexes in *N. benthamiana* leaves. We strongly recommend optimizing UV-crosslinking conditions according to tissue type, plastic film, and crosslinker combination. We provided the users with a proteomics dataset generated using this protocol, comprising six biological replicates. Two fractions were analyzed: the input and the OOPS interphase. Within the OOPS fraction, both crosslinked (CL) and non-crosslinked (No-CL) samples were processed, resulting in 18 samples analyzed. This experimental setup is essential for assessing RNA-binding protein enrichment.

**Figure 4:**
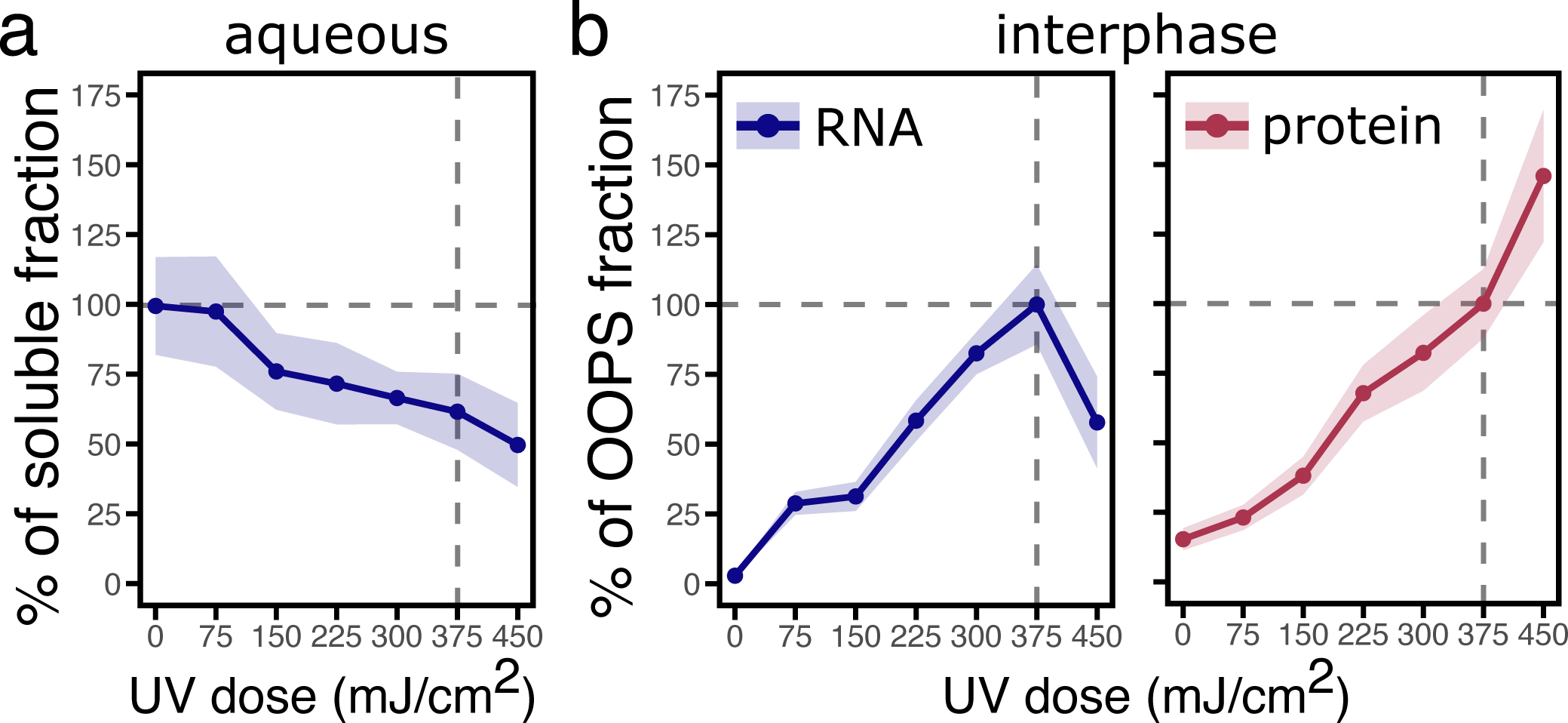
UV dose-response curve to determine optimal crosslinking conditions. Yields of RNA and protein were measured after irradiation with increasing UV doses. A. In the aqueous phase, RNA levels progressively decreased with increasing UV doses, likely due to crosslinking-induced relocation to the interphase. B. In the interphase, RNA and protein yields increased proportionally with UV irradiation. RNA levels in the aqueous phase of No-CL samples were considered 100%, while OOPS fractions were normalized to a dose of 375 mJ/cm^2^.

To ensure reproducibility, quality control analyses were conducted using Pearson correlation. As shown in Figure 5, the three sample groups clustered together, indicating high inter-replicate correlation. Additionally, we quantified the number of protein groups identified by LC-MS/MS. Consistent with previous results from or research group, the input fraction contained the highest number of protein groups, followed by the OOPS CL and OOPS No-CL fractions. To further explore the dataset, we performed principal component analysis (PCA). Here, the replicates of each sample clustered together, in agreement with the correlation analysis. Additionally, PCA revealed that the primary source of variability (PC1 > 67%) was attributable to UV crosslinking, followed by sample fractionation (PC2>21%). Collectively, these statistical analyses confirm that the protocol yields high-quality proteomics data. To identify the RNA-binding proteome (RBPome) of *N. benthamiana*, we performed enrichment analysis by comparing protein abundances between the OOPS CL and OOPS No-CL fractions. Proteins with a ≥2-fold increase (log2 FC>1) and adj. *p*>1.5 (-log10) were considered enriched. In Figure 6, a visual comparison of the datasets before and after filtering using these criteria is provided. After removing non-enriched proteins, a final set of 369 protein groups was retained. To benchmark the results, we examined the enrichment of bona fide RBPs, such as ribosomal proteins. A significant enrichment of ribosomal proteins in the OOPS CL fraction compared to the input (*p*<2.2e-16, Fisher’s exact test) validated the capacity of our protocol for isolating RBPs.

**Figure 5:**
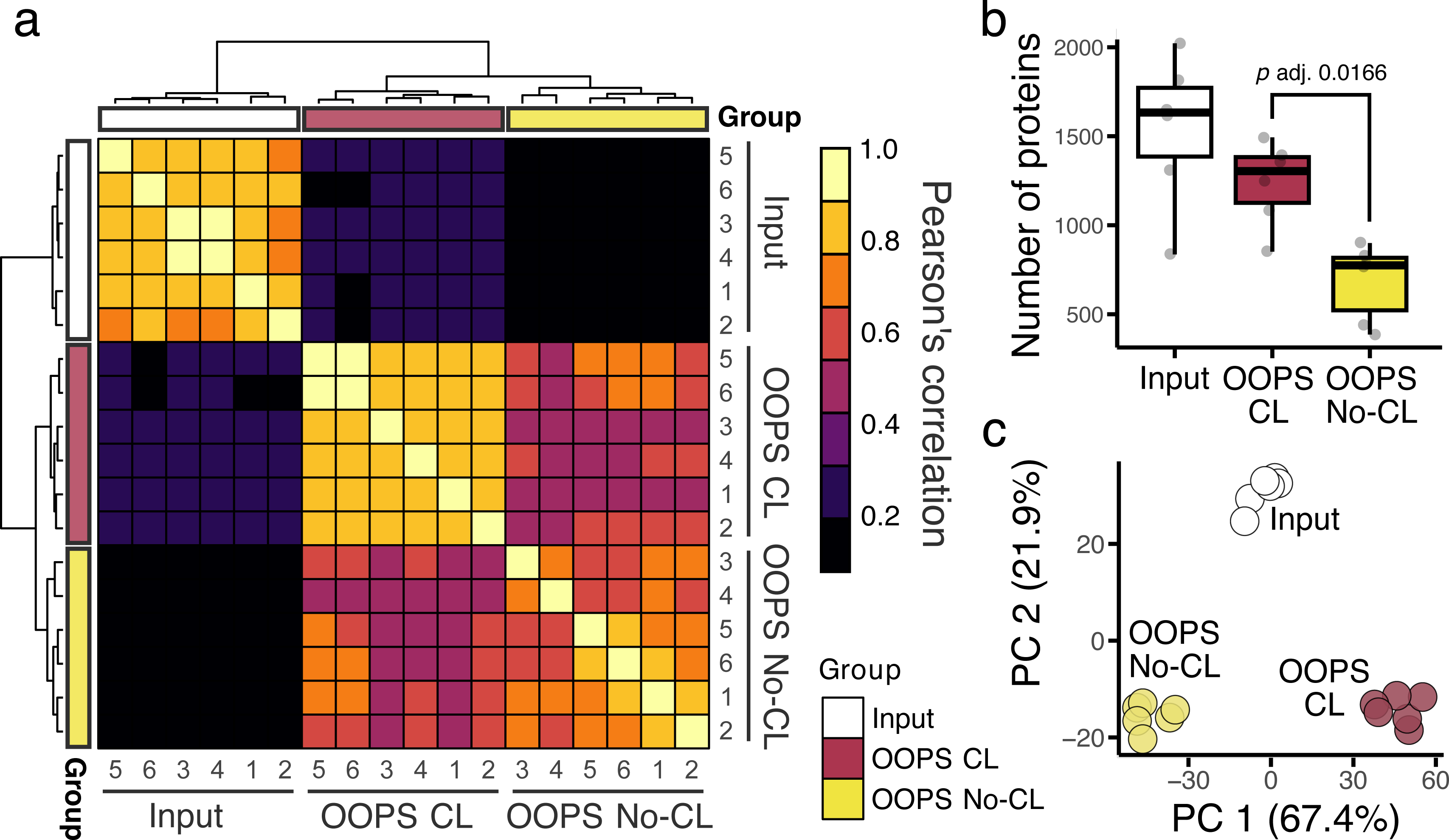
Quality control of proteomics data from OOPS fractions of *N. benthamiana* leaves. A. Pearson correlation analysis of label-free quantification (LFQ) values across six replicates of input, OOPS CL, and OOPS No-CL samples. Clustering of similar sample types is displayed along the axes. B. Protein group counts per fraction in the different samples. Significance was determined using Student’s *t*-test. C. Principal component analysis (PCA) reveals clustering of samples by fraction and crosslinking condition.

**Figure 6:**
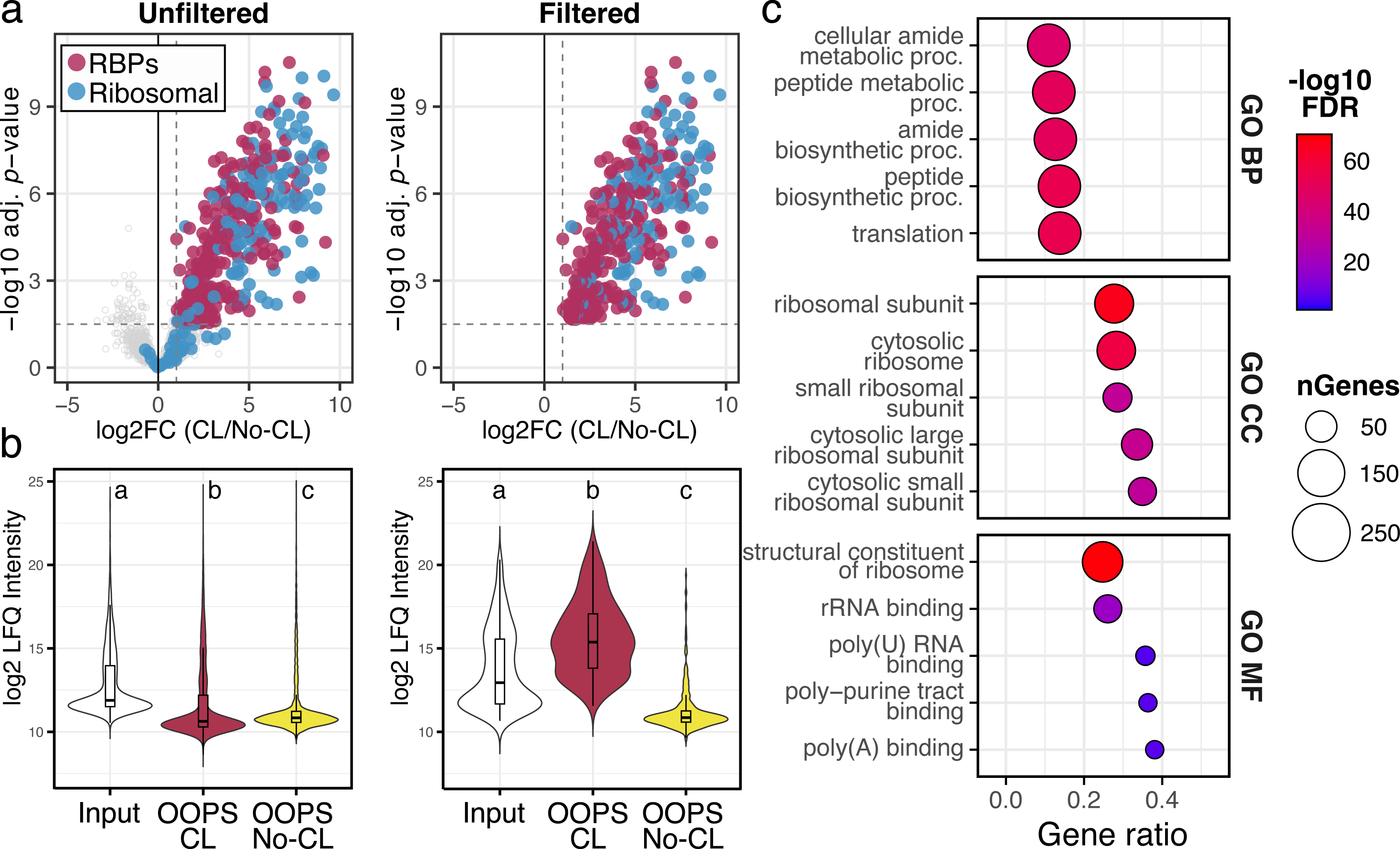
Enrichment analysis of RNA-binding proteins (RBPs) from *N. benthamiana* leaves. A. Volcano plots showing RBP enrichment in OOPS CL versus OOPS No-CL samples (log2 fold change ≥ 1), before and after data filtering. Ribosomal proteins (blue) and other RBPs (maroon are highlighted. Dashed lines indicate thresholds. B. Log2 LFQ values of proteins displayed in panel (A). Different letters indicate statistical differences (*p*<0.001, one-way ANOVA with Tukey’s *post hoc* test). C. Gene Ontology (GO) analysis of the filtered dataset showing the top five enriched terms per category: BP (biological process), CC (cellular component), and MF (molecular function). Color scale indicates -log10 FDR; bubble size, protein count; gene ratio, category representation.

We hypothesized that if a given RBP is crosslinked to RNA and subsequently captured in the interphase, its abundance would be higher in this fraction compared to total lysates. To test this, we compared LFQ values across fractions, before and after filtering. As expected, unfiltered datasets showed the highest abundance in the input fraction. However, in the filtered dataset of enriched RBPs, we observed significantly higher abundance in the OOPS CL fraction compared to both the input and OOPS No-CL fractions (mean differences = +1.94 and +4.43 log2 LFQ, respectively; adj. *p* < 0.001, one-way ANOVA with Tukey HSD *post hoc* test), supporting the successful enrichment of RNA-associated proteins.

Following dataset validation, we assessed whether RNA-related functional categories were enriched in the RBPome. Gene ontology analysis was conducted using ShinyGO (v0.82),^22^ revealing that the top five enriched terms in the categories of biological process, cellular component, and molecular function were associated with translation, ribosomal structures, and RNA binding, respectively. These results further confirmed the selective enrichment of RNA-binding proteins by our protocol.

Given that ribosomal proteins are highly expressed, we sought to rule out the possibility that their enrichment occurred by chance. We analyzed the top 10 enriched protein domains in the OOPS CL fraction and compared their distributions with the input fraction. In the OOPS CL fraction, domains related to ribosomal proteins, RNA recognition motifs (RRM), and S1 domains accounted for 40.4%, 23.3%, and 8.1% of proteins, respectively, compared to only 7.1%, 5.2%, and 1.8% in the input fraction (Figure 7). Overall, nine RNA-binding domains constituted over 80% of all motifs in the OOPS CL fraction but less than 25% in the input, indicating an enrichment of diverse RNA-associated domains and not only ribosome-related. In summary, this protocol effectively isolates the leaf RNA-binding proteome of *N. benthamiana*, generating high-quality datasets, potentially contributing to the advancement of RNA-protein interaction studies in plants.

**Figure 7:**
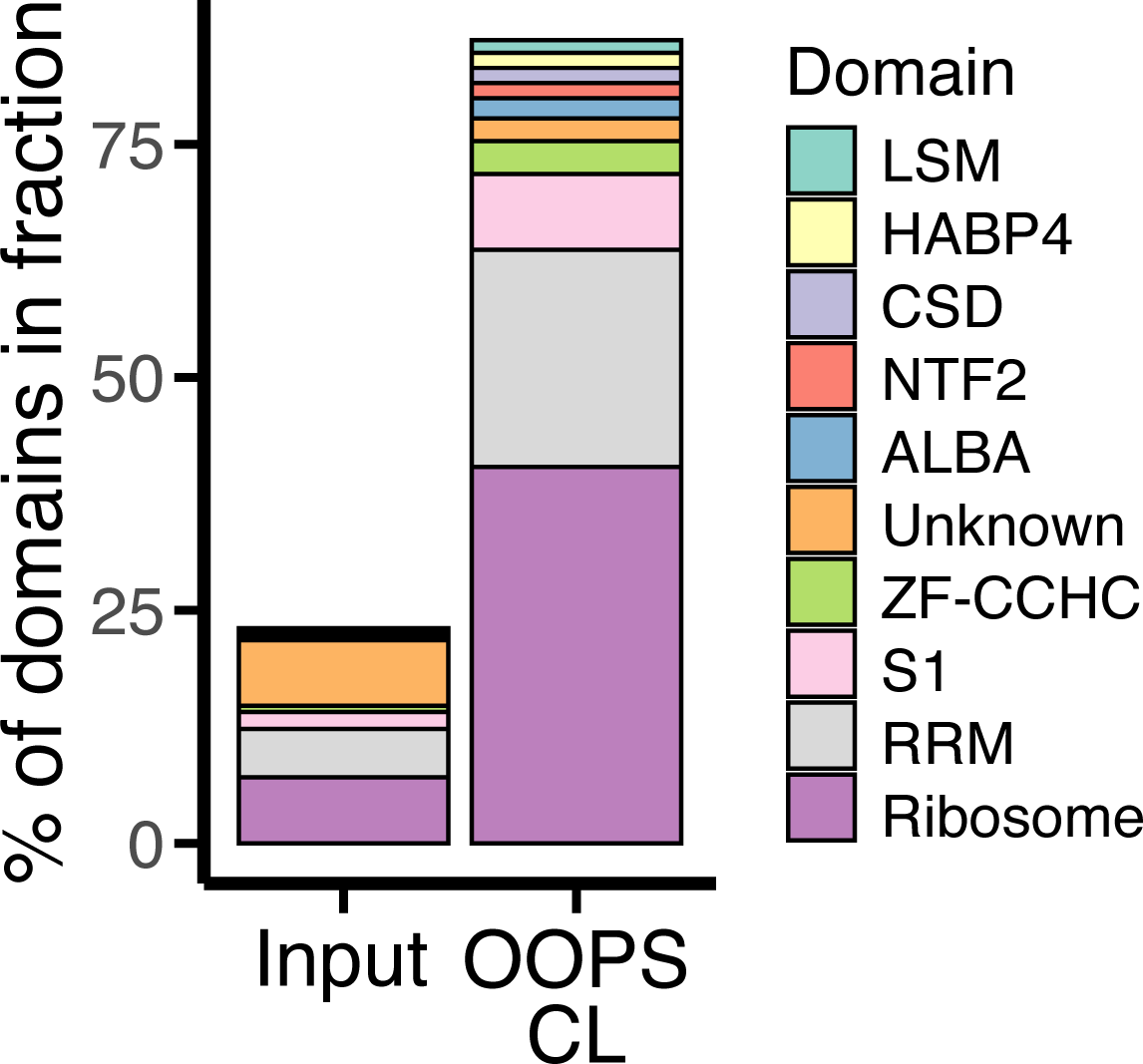
Protein domain analysis. Proportions of the top nine protein domains identified in the OOPS CL RBPome compared to their representation in the total leaf proteome. Unknown, indicates proteins lacking annotated domains in the Pfam database.^27^

### Limitations

This protocol was established using young *N. benthamiana* leaves, a relatively simple organ that lacks specialized modifications such as lignification and is characterized by its thin, flat structure. We acknowledge that adjustments may be required to optimize UV-crosslinking conditions for other tissues or species. One potential adaptation, as reported by others, is to perform UV irradiation on frozen, ground tissue.^23^

### Troubleshooting

Problem 1:

Low yield of RNA or RNA-binding proteins in the interphase.

Potential solution:

- Ensure that tissue crosslinking conditions (step 1) are optimized by performing a dose-response experiment, as exemplified in in Figure 4. Quantify RNA and protein yields in the aqueous phase and in the interphase to assess optimal crosslinking efficiency.
- Refer also to Troubleshooting 2, as incomplete lysis can compromise RNA/protein recovery.

Problem 2:

Incomplete tissue lysis.

Potential solution:

- If incomplete lysis is suspected, increase the tissue-to-lysis buffer ratio in step 11. The protocol uses a 1:1.5 (weight/volume) ratio. We have tested 1:2 and 1:2.5 ratios without compromising yield or lysis efficiency. However, increasing the buffer requires proportional adjustments of sodium acetate, phenol and chloroform volumes during phase separation in steps 16-17 (see main protocol).
- Alternatively, increase the shaking frequency and/or intensity during lysis incubation (step 12) to enhance mechanical disruption of the tissue.

Problem 3:

Poor compacting of the interphase during washing steps.

Potential solution:

- Increasing centrifugal force significantly improves the interphase integrity during re-extractions (steps 26-32). We routinely use 20,000 × g; however, the protocol showed to be effective when 15,000 × g was used.
- Extended incubations before phase removal can result in diffuse interphases. If this occurs, vortex the samples thoroughly and centrifuge again. To minimize this issue, reduce the number of samples processed in parallel to shorten handling times.
- If the issue persists despite repeat centrifugation, inefficient crosslinking may be the underlying cause (see Troubleshooting 1)

Problem 4:

Pellets containing RNA, proteins or RNA-protein complexes are difficult to resuspend (steps 39, 51, 63, 83 and 89).

Potential solution:

- Methanol and isopropanol precipitation lead to sample dehydration, making pellets difficult to dissolve. Complete removal of alcohol is essential for effective resuspension and downstream enzymatic treatments. After centrifugation:
- `o` Decant the supernatant and centrifuge briefly to collet residual droplets.
- `o` Use a pipette to remove any remaining alcohol.
- `o` Airdry the pellet.

**CRITICAL:** Avoid over-drying the pellet, as this may be not reversible. Based on experimental optimization, a drying period of 10 minutes in a flow cabinet is sufficient to eliminate ethanol without over-drying.

Problem 5:

Low number of proteins enriched in the OOPS CL fraction.

Potential solution:

- The protocol was optimized using 5 g of tissue input, with mass spectrometry performed using 150 µl of the resuspended interphase (equivalent to 2.5 g input; step 52a). The ratio between starting material and number of interphase re-extractions must be balanced to ensure sufficient RBP enrichment (see also Troubleshooting 6).
- To evaluate the protocol at lower input amounts, we tested 1 g of tissue and analyzed either i) half of the corresponding interphase (0.5 g equivalent), or ii) 0.5 g-equivalent interphase obtained from a 5 g input. We identified two key limitations in low-input conditions:
- `o` The RNase A/T1 concentration to release RBPs must be adjusted relative to the interphase amount. At standard conditions (2.5 g input and 1× RNase mix), RNase peptides comprised ∼20% of total iBAQ intensity (Figure 8). In contrast, for 1 g input with the equivalent RNase concentration, RNase accounted for up to 70%, limiting detection of enriched RBPs to only few hits.
- `o` LFQ intensity and total protein identifications decreased significantly (Figure 8). We hypothesize this is due to a molecular crowding effect mediated by RNA,^24^ which may enhance RNA-protein adducts partition and accumulation at the interphase. Reduced RNA concentrations in low-input samples may therefore impair RBP enrichment and detection.
- If low tissue input is required, we recommend scaling down RNase treatment proportionally.

**Figure 8:**
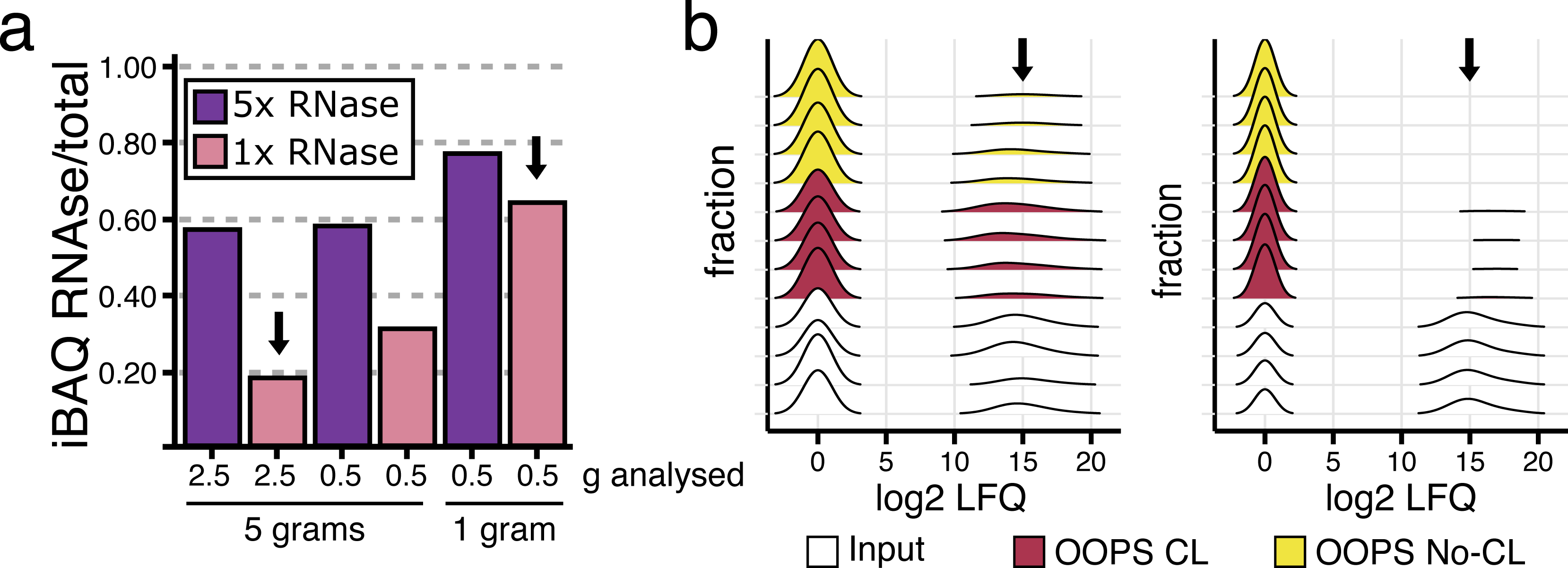
Optimization of tissue input and RNase ratio for proteomic analysis. A. LC-MS/MS analysis of OOPS CL samples from 5 g or 1 g of input tissue. Interphase fractions (equivalent to 2.5 g or 0.5 g) were treated with 1× or 5× RNase. Under optimal conditions (5 g, 1× RNase), RNase iBAQ signal was <30% while in 1 g samples, it exceeded 60%, compromising RBP detection. B. LFQ intensity distribution across fractions from four CL and four No-CL samples, prepared using 5 g (2.5 g interphase, left) or 1 g (0.5 g interphase, right). Input samples showed similar intensities for both experimental designs. However, LFQ values from the OOPS fractions using 5g were higher (indicated with arrows), consistent with the observations in panel (A). Peaks at zero correspond to imputed values.

Problem 6:

High proportion of non-RNA-associated proteins are identified in the crosslinked (CL) OOPS fraction.

Potential solution:

- This issue may arise from residual contamination due to incomplete removal of non-crosslinked proteins. We recommend increasing the number of re-extractions to improve interphase purity. Assess wash efficiency as follows.
- `o` Precipitate 300 µl of the organic fraction with methanol (steps 59 to 63) after each re-extraction and measure the protein concentration.
- `o` Normalize protein yield to the total volume of the organic fraction.
- `o` A progressive decline in protein content across washes is expected. Re-extraction is considered sufficient when protein yield is higher in the OOPS CL fraction than in the organic phase (see Figure 9).
- As an alternative, perform LC-MS/MS analysis of OOPS CL fractions after 2 to 6 re-extractions to determine the optimal number of washes.

**Figure 9:**
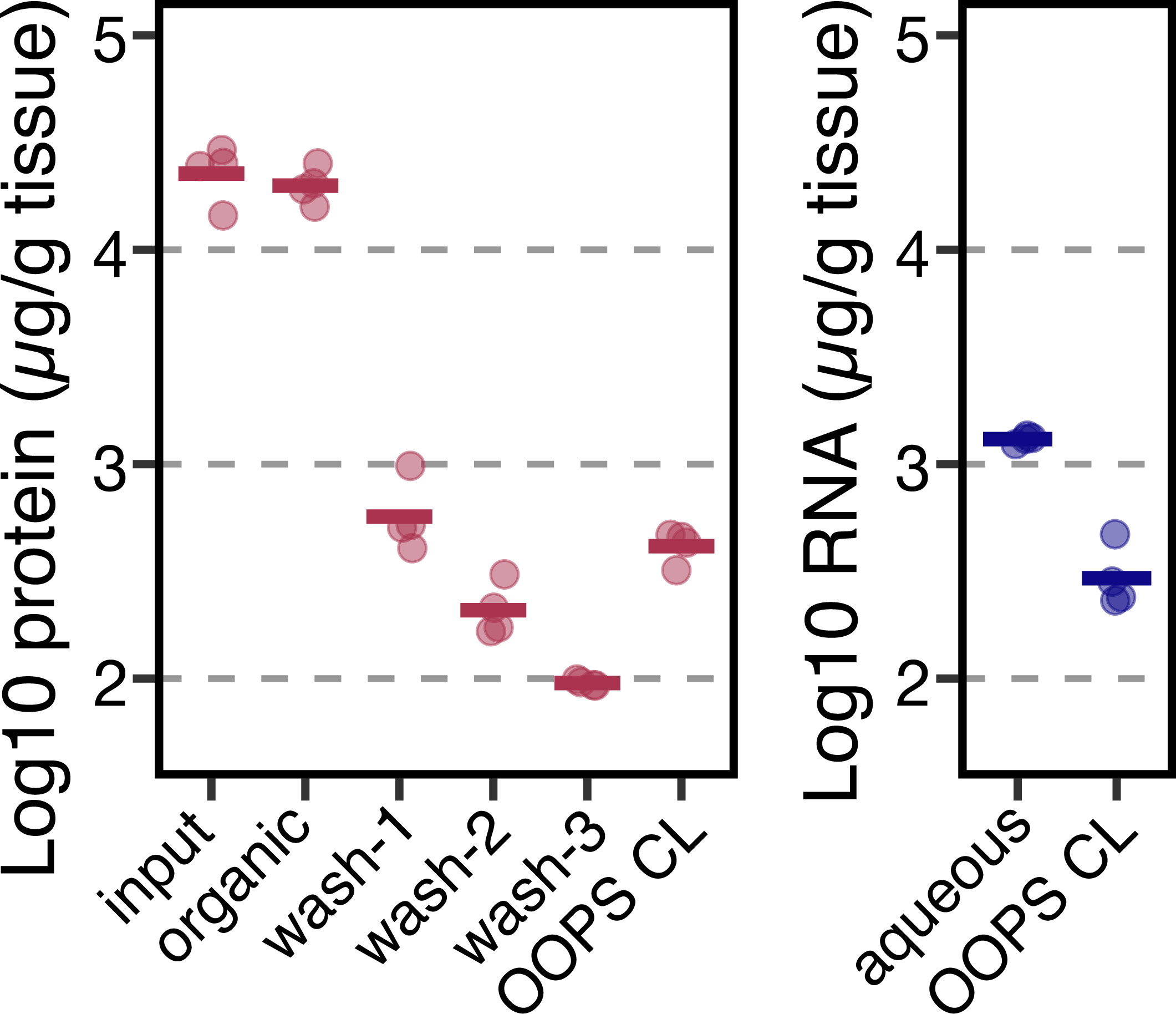
RNA and protein yield across the OOPS protocol. Protein and RNA were quantified throughout the protocol, from the initial lysate, organic phases during washes and the final OOPS CL fraction. Protein release from the interphase to the organic phase decreased progressively through washing steps, while the yields remained relatively high after three washes in the interphase (left panel). RNA yields in the final OOPS CL fraction were comparable to protein (right panel).

## Resource availability

### Lead contact

Further information and requests for resources and reagents should be directed to and will be fulfilled by the lead contact, Harrold van den Burg.

### Technical contact

Technical questions on executing this protocol should be directed to and will be answered by the technical contact, Victor A. Sánchez-Camargo.

### Materials availability

This study did not generate new unique reagents.

### Data and code availability

The datasets generated during this study are available at [name of repository]: [accession code/web link].

## Supporting information

supplemental_demo_data

## Acknowledgments

This research was supported by TKI T&U project (LWV20.105). We acknowledge Petra Houterman and Bas Beerens for technical assistance; and Harold Lemereis and Ludek Tikovsky for growing and caring the plants used in this research.

## Author contributions

V.A.S.C. designed and performed the experiments, analyzed and interpreted the data, and wrote the manuscript. G.K. contributed to the design of the proteomics experiments, performed LC-MS/MS sample analysis, participated in data interpretation, and reviewed the manuscript. H.A.B. conceived the research line, secured funding, contributed to experimental design and data interpretation, and critically reviewed the manuscript.

