## supplemental_demo_data for "Protocol for capturing the RNA-binding proteome from plants using orthogonal organic phase separation": volcano_rbpome.R

```
############################################################
## volcano_pair.R  –  Unfiltered vs. Filtered RBPome plots
## (with a single, in-panel legend on the right plot)
############################################################

## 0. Set working directory ---------------------------------------------------
setwd("/path/to/your/data")

## 1. Required packages -------------------------------------------------------
packages <- c("readr", "dplyr", "ggplot2", "ggrepel", "stringr", "patchwork")
invisible(lapply(packages, function(p) {
  if (!requireNamespace(p, quietly = TRUE)) install.packages(p)
}))
library(readr);  library(dplyr);  library(ggplot2)
library(ggrepel); library(stringr); library(patchwork)

## 2. File paths --------------------------------------------------------------
file_unf <- "starprotocol_rbpome.txt"          # unfiltered
file_flt <- "starprotocol_enriched_RBPome.txt" # filtered

## 3. Column names ------------------------------------------------------------
p_col     <- "-Log Welch's T-test p-value OOPS_XL_OOPS_NOXL"
fc_col    <- "Welch's T-test Difference OOPS_XL_OOPS_NOXL"
name_col  <- "Protein name"
ribo_col  <- "ribo_family"  # Column to highlight in maroon if filled

## 4. Helper functions --------------------------------------------------------
get_top_labels <- function(df, side = "right", n = 5) {
  if (side == "right") {
    df %>% filter(.data[[fc_col]] > 0) %>%
      arrange(desc(.data[[fc_col]])) %>% head(n)
  } else {
    df %>% filter(.data[[fc_col]] < 0) %>%
      arrange(.data[[fc_col]]) %>% head(n)
  }
}

wrap_labels <- function(x, width = 20) {
  stringr::str_wrap(x, width)
}

## 5. Read data ---------------------------------------------------------------
df_unf <- read_tsv(file_unf, show_col_types = FALSE)
df_flt <- read_tsv(file_flt, show_col_types = FALSE)

## 6. Prepare labels ----------------------------------------------------------
df_unf <- df_unf %>%
  mutate(!!name_col := str_split_fixed(.data[[name_col]], ";", 2)[, 1]) %>%
  mutate(label_wrap = wrap_labels(.data[[name_col]]))

df_flt <- df_flt %>%
  mutate(!!name_col := str_split_fixed(.data[[name_col]], ";", 2)[, 1]) %>%
  mutate(label_wrap = wrap_labels(.data[[name_col]]))

lab_unf_left  <- get_top_labels(df_unf, "left")
lab_unf_right <- get_top_labels(df_unf, "right")
lab_flt       <- df_flt %>% arrange(desc(.data[[fc_col]])) %>% head(5)

## 7. Volcano plot constructor ------------------------------------------------
make_volcano <- function(df,
                         labels_left  = NULL,
                         labels_right = NULL,
                         plot_title   = "",
                         show_legend  = FALSE) {
  
  df <- df %>%
    mutate(
      signif = (.data[[p_col]] > 1.5) & (.data[[fc_col]] > 1),
      ribo   = !is.na(.data[[ribo_col]]) & .data[[ribo_col]] != "",
      class  = dplyr::case_when(
        ribo   ~ "Ribosomal protein",
        signif ~ "RBP",
        TRUE   ~ "Other")
    )
  
  g <- ggplot(df, aes(x = .data[[fc_col]], y = .data[[p_col]])) +
    
    ## (i) background
    geom_point(
      data = subset(df, class == "Other"),
      colour = "lightgray", shape = 1, size = 1.5, alpha = 0.6,
      show.legend = FALSE
    ) +
    
    ## (ii) points entering the legend
    geom_point(
      data = subset(df, class != "Other"),
      aes(colour = class),
      shape = 16, size = 3.5, alpha = 0.85
    ) +
    
    ## colour scale & legend
    scale_colour_manual(
      values = c("RBP" = "maroon",
                 "Ribosomal protein" = "#4292c6"),
      labels = c("RBPs", "Ribosomal\nproteins"),
      guide = guide_legend(
        title = NULL,
        override.aes = list(size = 3)
      )
    ) +
    
    ## reference lines
    geom_vline(xintercept = 0,                        colour = "black") +
    geom_vline(xintercept = 1, linetype = "dashed",   colour = "grey50") +
    geom_hline(yintercept = 1.5, linetype = "dashed", colour = "grey50") +
    
    ## axes & theme
    scale_x_continuous("log2FC (CL/Non-CL)", limits = c(-5, 10)) +
    scale_y_continuous("–log10 adj. p-value",  limits = c(0, 11)) +
    ggtitle(plot_title) +
    theme_bw(base_size = 12) +
    theme(
      plot.title       = element_text(hjust = 0.5, face = "bold"),
      panel.grid.minor = element_blank(),
      legend.position  = if (show_legend) c(0.175, 0.89) else "none",
      legend.background = if (show_legend) element_rect(
        colour = "black",
        size = 0.4,
        fill = "white"
      ) else element_blank()
    )
  
  ## (iii) labels
  if (!is.null(labels_left)) {
    g <- g + geom_text_repel(
      data = labels_left,
      aes(label = label_wrap),
      max.overlaps = Inf, nudge_x = -0.25, size = 3, lineheight = 0.8
    )
  }
  if (!is.null(labels_right)) {
    g <- g + geom_text_repel(
      data = labels_right,
      aes(label = label_wrap),
      max.overlaps = Inf, nudge_x = 0.25, size = 3, lineheight = 0.8
    )
  }
  
  g
}

## 8. Build plots -------------------------------------------------------------
plt_unf <- make_volcano(df_unf,
                        labels_left  = lab_unf_left,
                        labels_right = lab_unf_right,
                        plot_title   = "Unfiltered",
                        show_legend  = FALSE)

plt_flt <- make_volcano(df_flt,
                        labels_right = lab_flt,
                        plot_title   = "Filtered",
                        show_legend  = TRUE)

## 9. Combine and display -----------------------------------------------------
combined <- plt_unf + plt_flt + plot_layout(ncol = 2, widths = c(1, 1))
#print(combined)

## 10. Optional save ----------------------------------------------------------
ggsave("volcano_rbps_starprotocol.pdf", combined, width = 12, height = 6)
```
