## Supplementary figures and images for "Protocol for capturing the RNA-binding proteome from plants using orthogonal organic phase separation"

### volcano_rbps_starprotocol.pdf

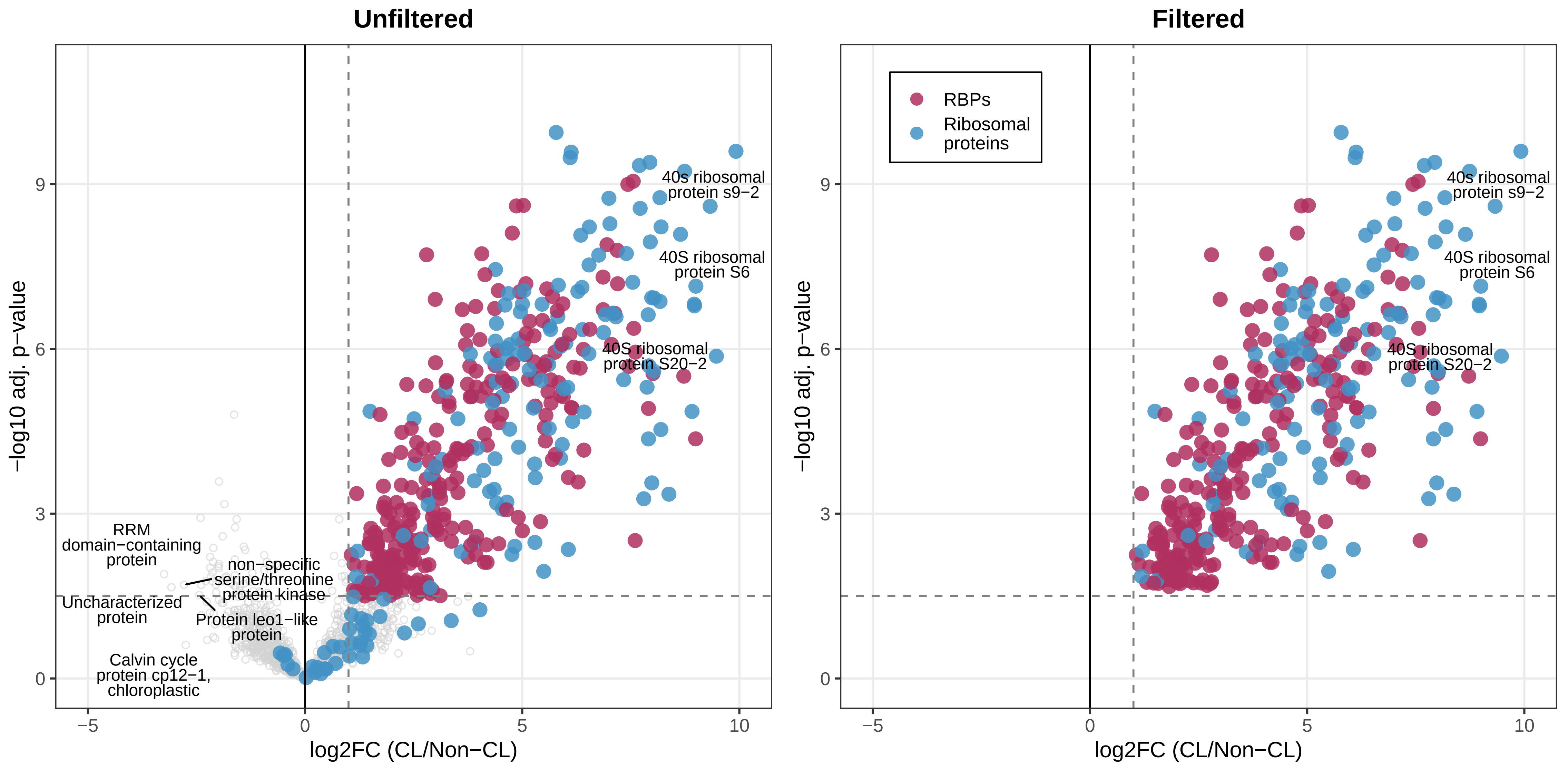
